# Sex-specific viability effects of mutations in *Drosophila melanogaster*

**DOI:** 10.1101/2024.03.09.584240

**Authors:** Robert H. Melde, JoHanna M. Abraham, Maryn R. Ugolini, Madison P. Castle, Molly M. Fjalstad, Daniela M. Blumstein, Nathaniel P. Sharp

## Abstract

In populations with separate sexes, genetic load due to deleterious mutations may be expressed differently in males and females. Evidence from insect models suggests that selection against mutations is stronger in males, with a positive intersexual correlation for fitness. This pattern will reduce deleterious allele frequencies at the expense of males, such that female mean fitness is greater than expected, preserving population persistence in the face of high mutation rates. While previous studies focus on reproductive success, mutation load depends on total selection in each sex, including selection for viability. In fruit flies, we might expect minimal sex differences in viability effects, since male and female larvae behave similarly, but many genes show sex-biased expression in larvae. We measured the sex-specific viability effects of nine “marker” mutations and 33 mutagenized chromosomes. We find that both types of mutations generally reduce viability in both sexes. Among marker mutations we detect instances of sex biased selection in both directions, but mutagenized chromosomes show little sign of sex-specific mutational variance. We conclude that some mutations can indeed affect viability in a sex-specific manner, but that the pattern of male-biased mutational effects observed previously for reproductive success is not apparent at the pre-reproductive stage.

## Introduction

Mutations are a foundational component of evolution and may threaten the persistence of species with high mutation rates (Anderson et al., 2004; Bull et al., 2007; Eigen, 1993; Gerrish et al., 2007, 2013; Matuszewski et al., 2017). Under classic mutation load theory (Haldane, 1937), a deleterious allele that arises at mutation rate *µ* will be held at frequency *q*\* ≅*µ*/*s* at mutation-selection balance, where *s* is the coefficient of selection against heterozygotes; the impact of such an allele on mean fitness in a diploid population is approximately 2*q*\**s* = 2*µ*, meaning that mutation load depends only on the mutation rate. Conceptually, mutation load is independent of the fitness effect because alleles that are more detrimental will be less frequent, severely affecting fewer individuals, and alleles with mild effects will be more frequent, slightly affecting more individuals. However, the fitness effects of mutations can potentially matter for the mutation load in populations with separate sexes.

An allele need not have the same effect on males as it does on females, i.e., *s*_*male*_ ≠ *s*_*female*_. This “sex-specific selection” could be sexually concordant, where *s*_*male*_ and *s*_*female*_ have the same sign but differ in magnitude, or sexually antagonistic, where *s*_*male*_ and *s*_*female*_ have opposing signs.

While the equilibrium frequency of a deleterious allele depends on the average selection coefficient across sexes, *s*_*avg*_, the fitness of mutant individuals depends on their sex. The mutation load experienced by females, which is most relevant for the absolute fitness of a population, is therefore 2*µ*(*s*_*female*_/*s*_*avg*_); compared with the classic prediction, sex-specific selection will reduce the mutation load experienced by females whenever *s*_*female*_/*s*_*avg*_ < 1, which occurs when *s*_*female*_ < *s*_*male*_ (Whitlock & Agrawal, 2009). In summary, sex-specific selection can enhance population productivity if deleterious mutations generally have stronger effects on males.

Instances of sex-specific selection acting through reproductive traits have been observed in both *Drosophila melanogaster* (Mallet et al., 2011; Mallet & Chippindale, 2011; Pischedda & Chippindale, 2005; Sharp & Agrawal, 2008, 2013, 2018; Sharp & Vincent, 2015; Whitlock & Bourguet, 2000) and the seed beetle *Callosobruchus maculatus* (Grieshop et al., 2016, 2021) where mutations tend to affect adult males more than adult females. This pattern is thought to arise primarily because of strong selection on male versus female reproductive traits, i.e., sexual selection. *Sexual selection* is commonly associated with *sex-specific selection* since traits that affect reproductive success are often sex-specific (e.g., sperm motility, sperm storage, etc.). When sexual and nonsexual selection act concordantly, i.e., alleles that increase male mating success are also beneficial for population mean fitness, sexual selection is thought to have a positive effect. Mutation load might also be reduced in sexual populations if there is positive assortative mating for genetic quality (Whitlock and Agrawal 2009), but empirical evidence for this type of non-random mating is mixed (Sharp & Agrawal, 2009; Sharp & Whitlock, 2019; Talagala et al., 2024). Finally, sexual selection creates a variety of costs that may outweigh the potential benefits (Bedhomme et al., 2008; Fedorka & Mousseau, 2004; Foerster et al., 2007; Gems & Riddle, 1996; Hotzy & Arnqvist, 2009; Mainguy et al., 2009; Partridge et al., 1986; Rowe & Rundle, 2021; Wigby & Chapman, 2005).

Importantly, sex-specific selection need not stem only from sexual selection, but the extent of sex-specific selection on traits unrelated to reproduction has not been well studied. In *Drosophila*, males and females display a number of differences in development and physiology, with females growing faster as larvae and achieving a larger adult body size (Millington & Rideout, 2018), and sex-specific quantitative trait variation is common (Li et al. 2024, Mackay & Anholt, 2006). Sex-biased gene expression has been identified in varying tissue types and life stages in numerous species (Grath & Parsch, 2016; Mank et al., 2010; Pauletto et al., 2018; Perry et al., 2014; Shi et al., 2016; Yasmin et al., 2022). Sex-specific selection may be mechanistically related to sex-biased gene expression: in *D. melanogaster*, sex-biased genes are associated with sex-biased fitness costs, where mutations tend to affect the fitness of whichever sex has greater expression (Connallon & Clark, 2011), and sex differences in human disease risk have also been linked to expression differences (Ober et al., 2008; Rubin et al., 2020). These studies suggest that sex-specific selection could act broadly in dioecious populations and may not be limited to reproductive traits. Sex-specific selection on juvenile viability, for example, could augment or counteract the benefits of sex-specific selection on adult reproductive success.

In this study, we tested for sex-specific effects of mutations on larval viability in *D. melanogaster*. We measured the relative viability of male and female larvae bearing one of nine visible mutant alleles, which tend to have large effects on fitness, or bearing one of 33 mutagenized chromosomes. We find that mutations generally reduce viability in both sexes, and that instances of sex-specific selection can occur for this trait, but that there is no significant bias towards one sex. We find little evidence for sex-specific mutational variance among mutagenized chromosomes. Based on a database of mutant phenotypes, we find a slight excess of genes with female-specific lethal alleles, which corresponds well with our measurements of mutagenized chromosomes. We conclude that individual mutations can have sex-biased viability effects, but without the male bias observed generally for reproductive traits.

## Methods

### General

We conducted our experiments using the Canton-S genetic background (hereafter referred to as wild-type, WT), which we maintain as a large outbred population, and reared flies at 25C with a 12h:12h light:dark cycle. All experimental flies were reared in standard vials on nutritionally defined yeast-sugar-agar medium supplemented with live yeast.

### Marker mutants

To test how specific mutations impact sex-specific larval viability, we measured the effects of nine marker alleles. Eight of these alleles are phenotypically dominant, one is phenotypically recessive, and all affect autosomal genes. The dominant alleles are *bw*^*D*^ (*brown-Dominant*), *Dr*^*1*^ (*Drop*), *wg*^*Gla-1*^ (*Glazed*), *Ki*^*1*^ (*Kinked*), *sens*^*Ly-1*^ (*Lyra*), *Sb*^*1*^ (*Stubble*), *sna*^*Sco*^ (*Scutoid*), and *U*^*2*^ (*Upturned*); we use abbreviated symbols for these alleles in the remainder of the text. In addition, we measured the effects of one recessive marker, *bw*^*1*^ (*brown*), in conjunction with the mutagenesis portion of this study (the markers *bw*^*D*^ and *bw*^*1*^ are different alleles of the same gene). We obtained these markers from the Bloomington Drosophila Stock Center and immediately back-crossed them into the wild-type genetic background for 8-10 generations. We maintained back-crossed dominant markers with balancer chromosomes prior to the experiments reported here; these markers have been shown to have deleterious effects on fitness components in prior studies (Civetta, 1999; Sharp & Agrawal, 2008; Tedman-Aucoin & Agrawal, 2012; Wang et al., 2009), but we are not aware of data on their sex-specific viability effects. We maintained the recessive *bw* allele in the homozygous state.

For each dominant marker, we began by crossing marked males from the stock population to WT virgin females in 20-30 vials. We then collected the resulting marked female offspring as virgins and crossed them to WT males in 20-30 vials. This provided one additional backcross generation just prior to our viability tests and removed any balancer chromosomes. We then collected the resulting marked males (heterozygous for the marker allele) and crossed them with WT virgin females, with two males and four females per vial (Fig. 1a). After four days of mating and oviposition, we discarded these adult flies. We scored the resulting offspring for both sex and marker status after 11 days of development, and again after a further 3 days. We scored offspring from one, two, or three separate blocks of crosses to ensure that many offspring were scored (>7500) for each marker allele. In the absence of any viability selection, we expect offspring to consist of equal numbers of marked females, marked males, WT females and WT males (Fig. 1a). We obtained an average of 102.4 scorable replicate vials for each marker (range 87-118) and scored a total of 69 681 flies in this part of the study.

**Figure 1.**
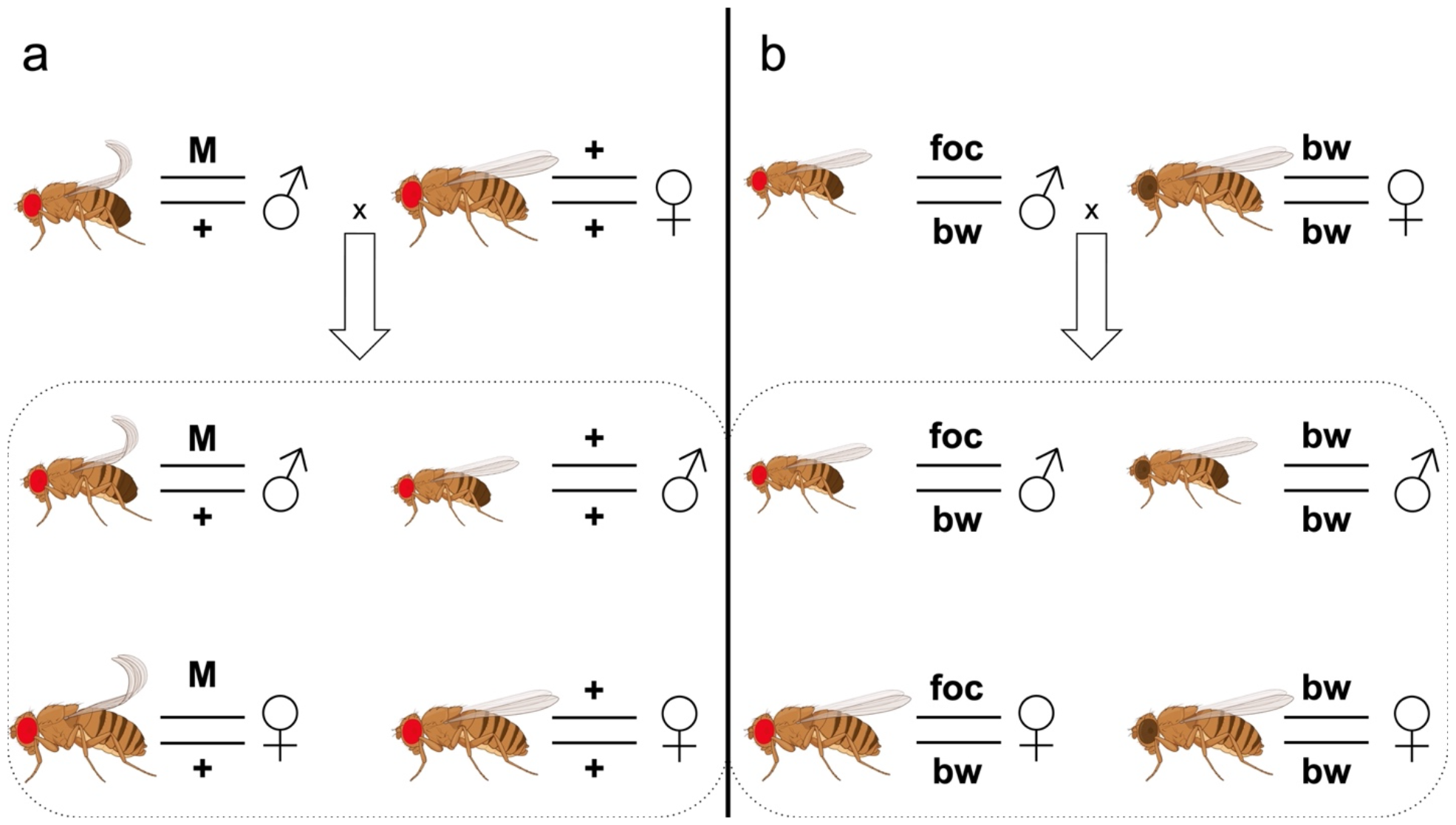
Crosses used to measure sex-specific larval viability. (a) We scored mutant (M) and non-mutant (+) males (♂) and females (♀) based on visible markers. Here we used single locus markers with dominant visible phenotypes located on chromosome 2 or 3. (b) We scored focal (foc/bw) and brown (bw/bw) males and females. Focal chromosomes either originated from control (non-mutagenized) or mutant (mutagenized) WT males.

### Mutagenesis

To test how a broader spectrum of mutations might impact sex-specific viability, we measured the effects of 33 unique sets of mutations created via a mutagenesis protocol (Fig S1). To create these, we held WT adult males without food and water in vials for 5 h prior to mutagen exposure. The mutagen solution included a 1:1 ratio of sucrose:water, a final concentration of 7.5 mM methyl methanesulfonate (MMS), and blue food coloring. Mutagen sensitivity can vary greatly across differing genetic backgrounds (Graf & Würgler, 1978; Snyder & Smith, 1982); previous reports use lower MMS concentrations, but our preliminary assays using 1.5 mM and 3 mM MMS did not show detectable viability effects in heterozygotes, and so we used 7.5 mM here. We exposed the starved adult males to the mutagen solution by saturating two pieces of filter paper and placing them in the bottom of fly bottles. The males were left to consume the solution for 24 hours, after which we selected males that had visible blue food coloring in their abdomen (indicating consumption of the mutagen). We allowed males to recover in standard food vials for about three hours prior to crossing. Each unique offspring from a mutagenized male is expected to carry a different set of mutations, and so we define each replicate “genotype” as stemming from a single male offspring of a mutagenized male. We expect mutagenesis to induce mutations on any chromosome; to standardize scoring across genotypes, we performed crosses to isolate only mutagenized second chromosomes, replacing the third chromosome, X chromosome and Y chromosome with un-mutagenized wild-type chromosomes (Fig. S1).

In tandem to this mutagenized treatment group, we performed equivalent crosses using males who were starved and fed a blue sucrose water solution without MMS. These acted as a control group and allowed us to infer the viability effects of mutagenized chromosomes separately from any effect of the *bw* marker. We measured the sex-specific viability of “focal” chromosomes, which were either mutagenized or un-mutagenized, in the heterozygous state, in competition with standard *bw*/*bw* flies (Fig 1b). In these crosses, offspring with red eyes carry the focal chromosome and offspring with brown eyes do not. In the absence of any viability selection, we expect offspring to consist of equal numbers of brown-eyed females, brown-eyed males, red-eyed females and red-eyed males. In total, we scored 20 158 flies in this part of the study: 8 757 from 100 vials in the control group, and 11 401 from 115 vials in the mutagenized group. The mutagenized replicates were established from 33 unique mutagenized genotypes.

The control replicates were sampled from a stock WT population, which is assumed to contain some genetic variation but for analysis purposes was considered as one “genotype”. This approach allows us to distinguish sex-specific effects of mutagenized chromosomes from sex-specific effects of the *bw* marker; we present the latter measures alongside our viability estimates for the dominant phenotypic marker alleles.

### Data Analysis

Let *k* be the total number of eggs of a given sex in a vial. In the absence of viability selection our crosses will result in equal numbers of wild-type and marked flies, with expected numbers *N*_*wt*_ = 0.5*k* and *N*_*mkr*_ = 0.5 *k w*_*mkr*_, respectively, where *w*_*mkr*_ represents the relative viability of the marker genotype. We can therefore calculate *w*_*mkr*_ = *N*_*mkr*_/*N*_*wt*_ for a given sex and marker. For comparison with other studies we can also calculate *s* = 1 – *w*_*mkr*_.

For the analysis of mutagenesis data, we need to consider both the effects of induced mutations and the effects of the *bw* marker used to identify the non-focal chromosome. In the control group where the focal chromosome is non-mutagenized, *N*_*wt,ctr*_/*N*_*mkr,ctr*_ equates to 1/*w*_*bw*_, where *w*_*bw*_ is the effect of the *bw* marker on viability in a given sex. In the treatment group where the focal chromosome is mutagenized, *N*_*wt,trt*_/*N*_*mkr, trt*_ equates to *w*_*mut*_/*w*_*bw*_, where *w*_*mut*_ is the effect of mutagenesis on viability in a given sex. We can therefore calculate *w*_*mut*_ = (*N*_*wt,trt*_/*N*_*mkr, trt*_)/(*N*_*wt,ctr*_/*N*_*mkr, ctr*_).

We analyzed our data and prepared our figures in *R* (v4.3.2), using the *lme4* package to fit generalized linear mixed-effect models (Bates et al., 2015). To analyze counts of marked and un-marked flies we fit binomial models (logit link), with a replicate-level random effect term to account for any overdispersion, and random effects of genotype and experimental block as appropriate. We tested for significant fixed effects using Wald tests and compared nested models using likelihood ratio tests (LRT). For each marker allele we first tested for an overall viability effect, i.e., a significant deviation of the mutant:non-mutant ratio from 0.5, regardless of sex. We then tested for a main effect of sex and sex-by-mutation interaction, retaining vial identity as a random effect. We report *P*-values with and without a Benjamini-Hochberg correction for multiple testing (Benjamini & Hochberg, 1995). We obtained mutant allele frequencies from the fixed effects of GLMMs and obtained 95% confidence intervals using parametric bootstrapping. Note that error bars are for visualization purposes and were not used to assess statistical significance.

For the mutagenesis data we first tested whether treatment group (mutagenized or non-mutagenized) had any effect on viability, regardless of sex; we take a significant effect of treatment as an indication that our mutagenesis procedure was successful. We also tested for a significant effect of genotype as an indication of mutational variance. We then tested for a treatment-by-sex interaction effect, which will indicate whether induced mutations, on average, have different effects on male versus female viability. We also considered the possibility that mutagenized chromosomes could display sex-biased effects in both directions, with no net bias when averaged across mutagenized genotypes. We tested for genetic variation in sex-specific effects in two ways. First, we tested for a random effect of genotype on the main effect of sex in a GLMM, using just the mutagenized group. Second, we defined θ_net_ as the log ratio of relative male viability to relative female viability for each replicate vial of each genotype and tested for a significant effect of genotype on θ_net_ using analysis of variance (ANOVA).

If any given mutation is equally likely to show a female-biased or male-biased effect on viability, and there are multiple viability-affecting alleles on each mutagenized chromosome, the variance of sex bias among chromosomes may underestimate the variance in sex bias among alleles. In other words, we may have limited power to detect genetic variation in sex biased effects (i.e., the kind of variation we observed among marker alleles) when female-biased and male-biased alleles are often found together on the same chromosome. To examine this possibility, we generated simulated datasets matching the structure of our mutagenesis data.

We simulated some number of viability-affecting mutations per genotype (Poisson distribution), ranging from 0.5 to 50 mutations per chromosome, each with some effect on sex-averaged viability (exponential distribution), holding the net effect of mutations on average viability constant, matching the effect of mutagenesis we observed (see Results). For each allele we simulated female- and male-specific effects based on a metric of sex bias, θ = *log*(*w*_male_/*w*_female_), with no average bias (E[θ] = 0), but with normally distributed variation among alleles, SD[θ] > 0. We considered scenarios with SD[θ] = 0.1, approximately the value observed for our marker mutant data, as well as higher values. We assume alleles have multiplicative effects on viability. Finally, we added among-replicate (error) variance of the same magnitude as our observed data and tested for significant genetic variance in θ_net_ using ANOVA as described above. For comparison, we used the same simulation model to examine power to detect a difference in the mean effect of mutations on males and females (E[θ] > 0).

### Database information

Given the thorough characterization of the *D. melanogaster* genome, we reasoned that a pattern of sex-biased viability effects might be apparent in the number of genes with known female- or male-specific lethal alleles. We used the QueryBuilder tool on *FlyBase*, release FB2023_06 (Gramates et al., 2022), to screen for alleles with a phenotypic description containing the terms “lethal”, and “male” or “female”, considering only genes with a “current” gene model status. This will include alleles annotated with similar terms, e.g., “partially lethal”. We further screened this dataset by discarding alleles described as having lethal effects in both sexes. Finally, we obtained a list of unique genes affected by these alleles, and the coding sequence length of those genes.

## Results

### Marker alleles

Sex-specific viability data for eight dominant markers are given in Supplementary Information, Dataset 1; data for the recessive *bw* marker are given in Supplementary Information, Dataset 2. Pooling data for males and females, we detected significant variation in viability effects across the nine markers (LRT for marker effect: χ^2^ = 91.13; *df* = 8, *P* = 2.75 × 10^−16^). Considering each marker separately, we found that *Gla, Ki, Ly, Sco, U*, and *bw* significantly decreased larval viability, *bw*^*D*^ significantly increased larval viability, and *Dr* and *Sb* had no significant effect on larval viability (Table 1). Our estimates of viability effects are positively correlated with published estimates for the same alleles on other genetic backgrounds, indicating that these measures are repeatable (Sharp & Agrawal, 2008; Wang et al., 2009).

**Table 1:** Estimates of larval viability effects of visible marker alleles.

| Marker | Flies scored | Vials | Non-sex-specific viability |  |  | Sex-specific viability |  |  | Selection coefficients |  |
| --- | --- | --- | --- | --- | --- | --- | --- | --- | --- | --- |
|  |  |  | *z* | *P* | Adj. *P* | *z* | *P* | Adj. *P* | *S<sub>female</sub>* | *S<sub>male</sub>* |
| *bw<sup>D</sup>* | 11047 | 118 | 2.88 | $1.49 \times 10^{-3}$ | $2.23 \times 10^{-3}$ | 0.73 | 0.46 | 0.59 | -0.05 | -0.08 |
| *Dr* | 9890 | 99 | -1.26 | 0.21 | 0.23 | -0.13 | 0.90 | 0.92 | 0.03 | 0.03 |
| *Gla* | 8499 | 87 | -3.98 | $7.01 \times 10^{-5}$ | $1.50 \times 10^{-4}$ | 2.20 | $2.76 \times 10^{-2}$ | $8.28 \times 10^{-2}$ | 0.17 | 0.08 |
| *Ki* | 7819 | 100 | -3.35 | $8.16 \times 10^{-4}$ | $1.47 \times 10^{-3}$ | 2.83 | $3.33 \times 10^{-3}$ | $1.50 \times 10^{-2}$ | 0.19 | 0.07 |
| *Ly* | 8694 | 99 | -9.53 | $< 2 \times 10^{-16}$ | $9.00 \times 10^{-16}$ | 0.99 | 0.32 | 0.58 | 0.21 | 0.18 |
| *Sb* | 8168 | 113 | -0.93 | 0.35 | 0.35 | -1.80 | $7.27 \times 10^{-2}$ | 0.16 | -0.02 | 0.06 |
| *Sco* | 7890 | 101 | -10.20 | $< 2 \times 10^{-16}$ | $9.00 \times 10^{-12}$ | 0.10 | 0.92 | 0.92 | 0.24 | 0.23 |
| *U* | 7674 | 102 | -6.52 | $7.21 \times 10^{-11}$ | $2.16 \times 10^{-10}$ | -0.86 | 0.39 | 0.59 | 0.16 | 0.20 |
| *bw* | 8757 | 100* | -2.46 | $1.38 \times 10^{-2}$ | $1.77 \times 10^{-2}$ | -4.44 | $8.86 \times 10^{-6}$ | $7.97 \times 10^{-5}$ | 0.04 | 0.21 |

\*An additional 100 vials were used for strains with mutagenized chromosomes.

Our main interest was whether alleles have sex-specific viability effects. Considering all nine marker alleles, we found strong evidence that the effect of sex on the viability of marked flies varied among markers (LRT for sex-by-marker interaction: χ^2^ = 39.64; *df* = 8, *P* = 3.74 × 10^−6^). Considering each marker separately, we found that three showed evidence for sex-specific larval viability effects (Table 1, Figure 2): for *Ki* and *Gla* the marker was more deleterious in females, whereas the recessive allele *bw* was more deleterious in males. The significant effect of sex on the viability effect of *Gla* did not survive correction for multiple testing (Table 1).

**Figure 2.**
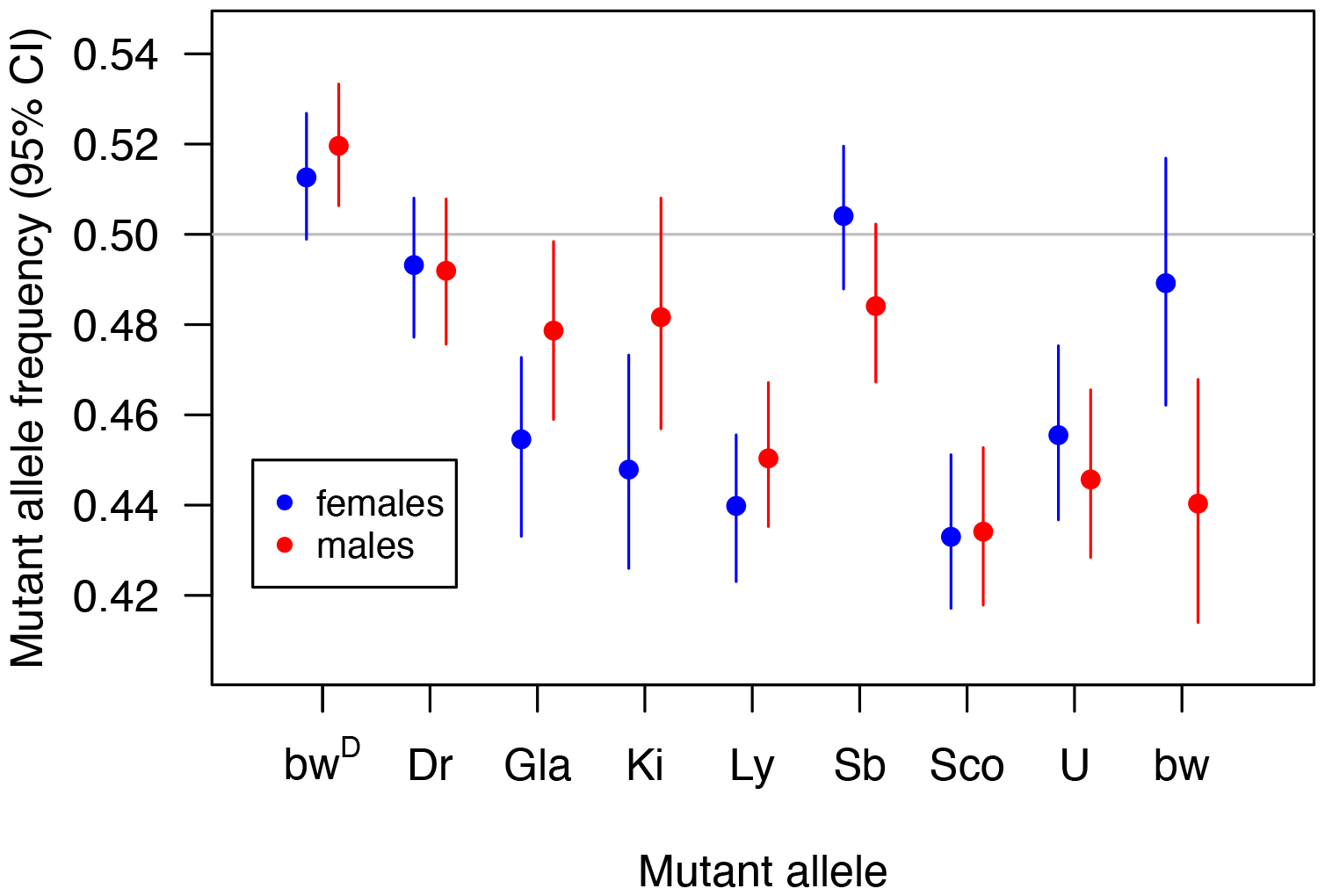
Observed mean frequencies of marker mutation bearing offspring for eight different phenotypically dominant markers. Offspring were either mutant or wild type and were scored in a sex-specific manner to calculate observed frequencies seen here. Averaged across sexes, *Gla, Ly, Ki, Sco*, and *U* significantly decreased larval viability, *bwD* significantly increased larval viability, and *Dr* and *Sb* had no significant effect on larval viability. *Gla* and *Ki* show evidence for sex-specific larval effects where females are more strongly affected, whereas males were more strongly affected by the *bw* allele. Confidence intervals were obtained using parametric bootstrapping.

We found that the viability selection coefficients (Table 1) of the nine marker alleles showed a positive intersexual correlation (Spearman’s rank correlation: *ρ*= 0.70, *P* = 0.043) indicating that the effects of marker alleles on viability are generally sexually concordant (Figure 3).

**Figure 3.**
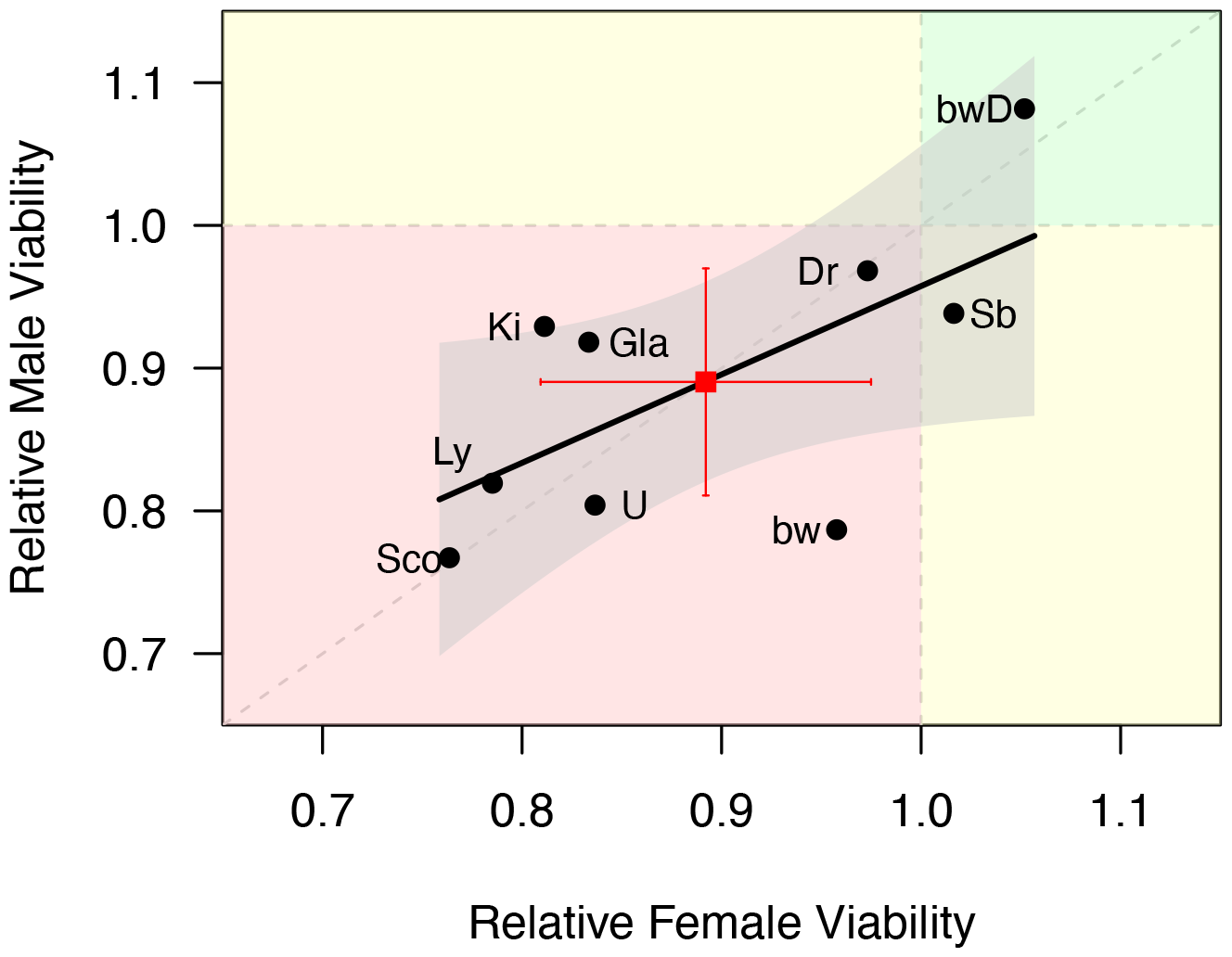
Sex-specific viability of marker mutation bearing chromosomes compared to wild type chromosomes. Points in quadrant one show increased viability for both males and females (green). Points in quadrant two show viability increased in males and decreased in females (yellow). Points in quadrant three show viability decreased in both males and females (red). Points in quadrant four show viability increased in females and decreased in males (yellow). The shaded region represents a 95% confidence band for the line of best fit from a linear regression model.

### Mutagenesis

Sex-specific viability data for 33 mutagenized chromosomes and corresponding controls are given in Supplementary Information, Dataset 2. Before considering sex-specific effects of mutagenized chromosomes, we tested whether our mutagenesis protocol produced a measurable effect on viability in general. We found that treatment group had a significant effect on viability (*z* = –3.67, *P* = 2.41 × 10^−4^) where undergoing mutagenesis, on average, decreased survival odds by 0.22. Comparing this effect to the mutation accumulation results from Sharp and Agrawal (2018) for the viability of heterozygous second chromosomes, our mutagenesis was equivalent to 61 generations of spontaneous mutation accumulation. We also detect significant genetic variation for viability among mutagenized genotypes (LRT: χ^2^ = 7.40; *df* = 1, *P* = 6.51 × 10^−3^); this mutational variance is equivalent to 75 generations of mutation accumulation (Sharp and Agrawal 2018). These findings confirm that our mutagenesis protocol induced a sufficient set of mutations in order to generate detectable changes in the mean and genetic variance of larval viability.

We found a significant positive intersexual correlation for relative larval viability (*r* = 0.57, *df* = 31, *P* = 5.78 × 10^−4^) and observed that most genotypes showed reduced relative viability in both sexes (Figure 4), indicating that mutations typically reduce viability in a sexually-concordant fashion.

**Figure 4.**
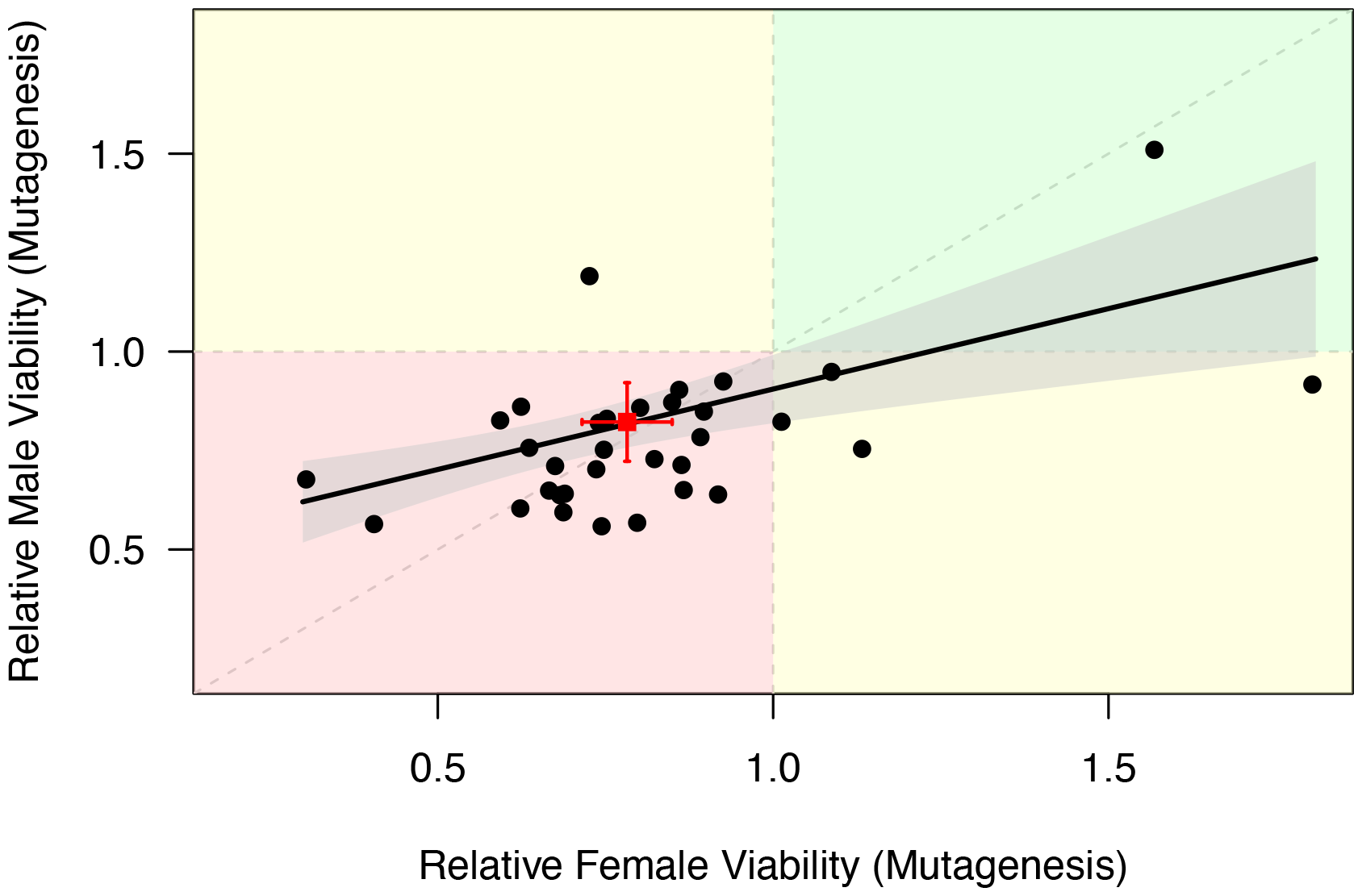
Relative viability for males and females of 33 genotypes bearing randomly induced mutations. Points in quadrant one show increased viability for both males and females (green). Points in quadrant two show viability increased in males and decreased in females (yellow). Points in quadrant three show viability decreased in both males and females (red). Points in quadrant four show viability increased in females and decreased in males (yellow). The shaded region represents a 95% confidence band for the line of best fit from a linear regression model.

To test for sex-specific viability effects, we ran a GLMM with the inclusion of a sex-by-treatment interaction term. This term is important since we used *bw* as a marker to indicate the absence of the focal chromosome. If we do not take this into account, any sex-specific effects of *bw* would be conflated with sex-specific effects of mutagenesis. We found that mutagenesis reduced viability to a greater extent in females than in males (a decrease in survival odds of 0.24 versus 0.20, or a 1.2-fold greater effect on females), but this difference was not statistically significant (*z* = 0.96; *P* = 0.34). In other words, we did not see evidence of a sex bias in the average effect of induced mutations on viability. This does not preclude the possibility that mutant genotypes could have sex-specific effects, but with a similar frequency of female biased and male biased mutant genotypes. However, we did not find a significant random effect of genotype on the main effect of sex (LRT: χ^2^ = 0.24, *df* = 1, *P* = 0.62), nor did we find significant genetic variance for θ, a metric of the sex difference in relative viability calculated for each replicate (ANOVA: *F* = 2.31; *df* = 1; *P* = 0.13).

In summary, we did not detect significant sex-specific viability effects of induced mutations, either in terms of a general bias towards a given sex, or in terms of variability in sex differences among mutant genotypes. We performed simulations to evaluate the statistical power that an assay of the kind we performed would have to detect differences in the mean or genetic variance of sex differences. In the presence of a net bias of mutational effects towards one sex or the other, we find that detection power is very high, regardless of whether fitness decline is due to few mutations of large effect or many mutations of small effect (Fig. 5A). By contrast, our simulations suggest that the power to detect variance in sex bias among alleles, based on their aggregate effects on each genotype, is modest at best, not exceeding 40% (Fig. 5B).

**Figure 5.**
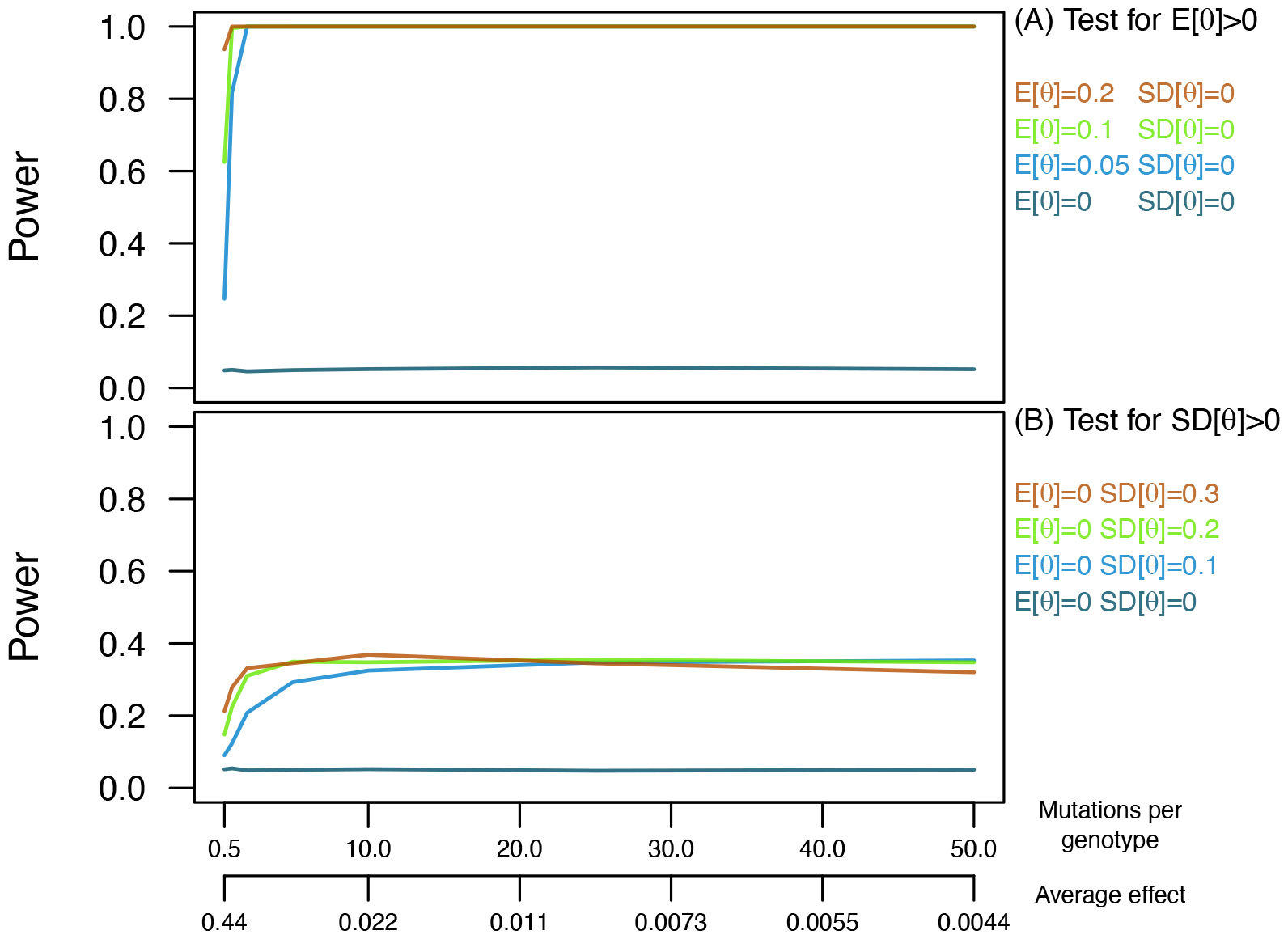
Power to detect nonzero mean or variance of sex-specific viability effects. We simulated datasets with the same structure as our mutagenized genotypes, maintaining the same overall impact on sex-averaged fitness, ranging from few mutations of large effect (towards the left) or many mutations of small effect (towards the right). (A) An assay of the kind we performed would have very good power to detect a difference in the average effect of mutations on male versus female viability (E[θ] > 0). (B) An assay of the kind we performed would have limited power to detect variance in sex biased effects among alleles (SD[θ] > 0) when there is no sex bias on average. Our measures of sex-specific effects of marker alleles would suggest a value of SD[θ] ≈ 0.1. In the absence of any effects (E[θ] = 0, SD[θ] = 0), both types of test are significant about 5% of the time, as expected.

### Database information

Data obtained from FlyBase regarding alleles with sex-specific lethal effects are given in Supplementary Information, Dataset 3. We found 392 alleles described as having lethal effects on just one sex. Of these, 242 (61.7%) are female-specific lethals. The interests of the research community could drive this pattern if, for example, genes with female-specific effects are studied more intensively through the generation of more mutant alleles. We might expect less bias in the number of genes known to be affected by such alleles; we find 298 genes known to have sex-specific lethal alleles: 161 in females, 130 in males, and 7 genes where both female-specific and male-specific lethal alleles have been described. These results suggest that 1.24 times as many genes are potentially subject to female-specific lethal mutations than male-specific lethal mutations. Accounting for the coding sequence length of these genes, this ratio becomes 1.28. We note a similarity between these results and the observation above that the effect of mutagenesis on viability is 1.2-fold greater in females.

## Discussion

The total effect of mutations on each sex is key to predicting how the mutation rate will affect population productivity (Whitlock & Agrawal, 2009). There is good evidence that many mutations reduce reproductive success in both sexes, but with a more severe effect on males (Grieshop et al., 2016, 2021; Mallet et al., 2011; Mallet & Chippindale, 2011; Pischedda & Chippindale, 2005; Sharp & Agrawal, 2008, 2013, 2018; Sharp & Vincent, 2015; Whitlock & Bourguet, 2000). Our goal was to assess the extent to which mutations have sex-specific effects on another key component of fitness––viability. Among studies of mutational effects, egg-to-adult viability in *D. melanogaster* is a commonly measured fitness component. As in prior studies, we see that mutations often reduce viability and rarely increase it, but we took the additional step of measuring the viability of males and females separately. Among nine “marker” alleles with visible phenotypic consequences, we detected three cases of sex-specific effects; of the two cases that survive correction for multiple testing, one is more harmful to males and the other is more harmful to females (Fig. 2). Among 33 chromosomes carrying induced mutations in the heterozygous state, we do not find evidence that mutations affect viability in one sex more than the other (Fig. 4); an experiment of this size would likely have high power to detect such a difference (Fig. 5A).

Although we found significant genetic variance in the sex-averaged effects of mutagenized chromosomes, we did not detect significant genetic variance in the degree of sex bias. At face value, this is at odds with our findings for marker mutations, which suggest that mutations with male-biased and female-biased viability effects may occur with similar frequency and are not too rare. However, power simulations indicate that genetic variance in sex bias may be hard to detect when mutations with opposing effects are aggregated on mutagenized chromosomes (Fig. 5B). Taken together, our experimental results therefore support the existence of deleterious alleles with sex-specific effects on viability, while also indicating that there is little to no bias towards one sex or the other in terms of average severity.

In terms of mutation load, the degree of sex differences in viability effects will only be relevant if the impact of mutations on viability is not very small relative to the impact of mutations on adult fitness. Previous studies show that the effect of an allele or mutant chromosome on viability represents 30-60% of its combined effect on viability and adult reproductive success (e.g., marker alleles: Sharp & Agrawal (2008); homozygous mutation accumulation chromosomes: Sharp & Agrawal (2013); heterozygous mutation accumulation chromosomes: Sharp & Agrawal (2018)), meaning that viability should ideally not be ignored when considering sex-specific mutational effects. Five of the marker alleles used in this study were included in the Sharp & Agrawal (2008) study, which measured male siring success, female egg production, and sex-averaged larval viability. Here we used a different genetic background, and we find weaker overall viability effects (paired t-test: *t* = 3.45, *df* = 4, *P* = 0.026). At face value, combining the results of these experiments shows cases where the extent of male-biased selection inferred from reproductive success would be counteracted (*Gla, Ly*), amplified (*Sb*), or little changed (*Dr*) by the incorporation of sex-specific viability selection.

While our data support the notion that most mutations are deleterious, we found that the allele *bw*^*D*^ increased viability in both sexes, in contrast to the *bw*^*1*^ allele (Fig. 2). Using a different genetic background, Wang et al. (2009) found that *bw*^*D*^ increased sex-averaged viability in both of two media environments. This allele is believed to result from the spontaneous insertion of a large heterochromatic sequence into the *bw* coding region (Suso Platero et al., 1998). While *bw*^*D*^ may be detrimental to other fitness components, the repeated finding that this allele enhances viability may warrant further investigation.

Our analysis of information from *FlyBase* reinforces our conclusion that alleles with sex-specific viability effects are not rare. The effects of mutant alleles on fitness components are generally not measured and catalogued in *FlyBase*, but recessive lethality represents an exception. We found that hundreds of genes have alleles with sex-specific lethal effects, with about 1.2 times as many genes associated with female-specific lethals as male-specific lethals. These data do not necessarily represent a random sample of alleles, but we think it unlikely that researchers are biased towards reporting lethality in one sex over the other. Interestingly, point estimates from our mutagenesis data also indicate an average effect on female viability that is about 1.2-times the average effect on male viability, suggesting that sex-specific lethals may reflect a more general pattern of sex-biased fitness effects. In a study of associations between standing genetic variants and adult fitness in *D. melanogaster*, in which adult survival was potentially a large component of fitness, Singh et al. (2023) found evidence for stronger overall selection on females. Taken together, these findings illustrate that sex differences in selection are likely more complex than expected under sexual selection alone.

## Acknowledgements

Stocks obtained from the Bloomington Drosophila Stock Center (NIH P40OD018537) were used in this study. Thanks to E. Anderson, C. Dong, J. Hintz, B. Hoyt-Glennon, N. Lacke, M. Lamps, T. Lee, J. Pfeifer, N. Prudlo, A. Smith, C. Smyth, L. Torhorst, and Y. Wang for assistance in the lab.

## Supplementary Information

**Supplemental Figure 1.**
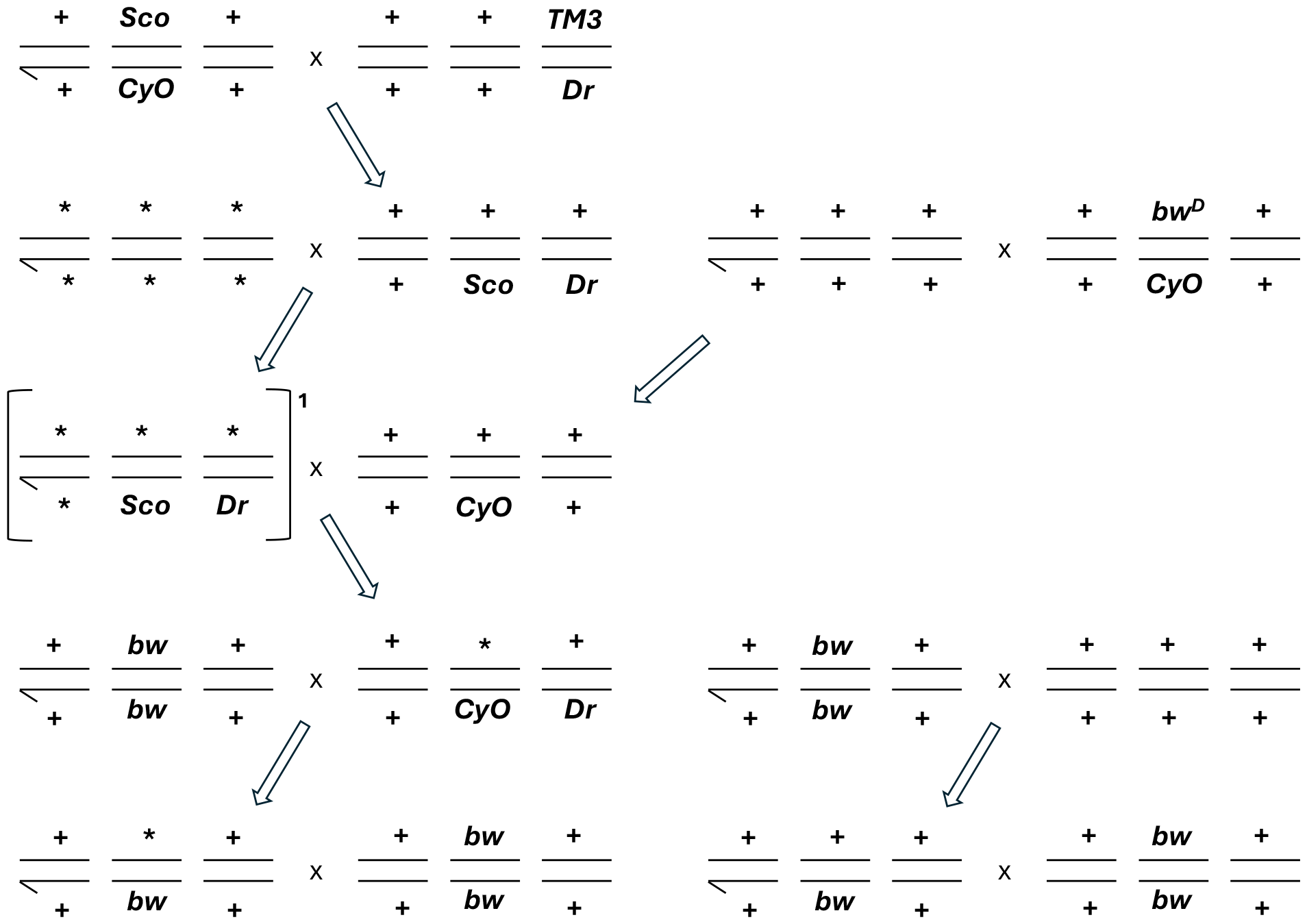
Crossing scheme used to prepare mutagenized strains. The purpose of these crosses was to situate mutagenized second chromosomes on a standard genetic background for viability assays. Lines represent chromosomes 1–3. Chromosomes denoted with * originate from males that consumed MMS, and those denoted with + are wild type. Additional dominant markers (*bw*^*D*^, *Sco, Dr*) and balancer chromosomes (*CyO, TM3*) were used to but were selectively removed prior to the assay. All crosses involved virgin females. At the third step, single-male bottlenecks (square brackets) were used to establish independent “genotypes” with induced mutations. 50 of these bottlenecks were performed, from which 33 scorable genotypes were successfully propagated.

## Notes

### Competing Interest Statement

The authors have declared no competing interest.

